# Human telomere repeat binding factor TRF1 replaces TRF2 bound to shelterin core hub TIN2 when TPP1 is absent

**DOI:** 10.1101/436014

**Authors:** Tomáš Janovič, Martin Stojaspal, Pavel Veverka, Denisa Horáková, Ctirad Hofr

## Abstract

Human telomeric repeat binding factors TRF1, TRF2 along with TIN2 form a core of shelterin complex that protects chromosome ends against unwanted end-joining and DNA repair. We applied a single-molecule approach to assess TRF1-TIN2-TRF2 complex formation in solution at physiological conditions. Fluorescence Cross-Correlation Spectroscopy (FCCS) was used to describe the complex formation by analyzing how coincident fluctuations of differently labeled TRF1 and TRF2 correlate when they move together through the confocal volume of the microscope. We observed, at the single-molecule level, that TRF1 effectively substituted TRF2 on TIN2. We assessed the effect of another telomeric factor TPP1 that recruits telomerase to telomeres. We found that TPP1 upon binding to TIN2 induces allosteric changes that expand TIN2 binding capacity, such that TIN2 can accommodate both TRF1 and TRF2 simultaneously. We suggest a molecular model that explains why TPP1 is essential for the stable formation of TRF1-TIN2-TRF2 core complex.

## Introduction

Human telomeres are maintained by telomerase^1,2^ and protected by telomeric proteins^3,4^. Telomeric proteins recruit telomerase to telomeric DNA^5^. Shelterin is a six-protein complex comprising of TRF1, TRF2, TIN2, TPP1, POT1 and RAP1. Shelterin associates specifically with telomeric DNA repeats and protects linear chromosome ends from being recognized by the DNA repair machinery as damaged DNA^4^. TRF1 and TRF2 bind the double-stranded telomeric DNA^6,7^. TRF2 protects chromosome ends mainly by invasion of the 3’ single-stranded overhang into the duplex telomeric repeats, thus forming protective lasso-like structures known as telomeric loops^8,9^. RAP1 interacts solely with TRF2 and regulates specificity of TRF2’s binding to telomeric DNA and subsequent telomeric loop processing by helicases^10,11^.

TIN2 (TRF1-Interacting Nuclear factor 2)^12^ binds both factors, TRF1 and TRF2 (Telomere Repeat-binding Factor 1 and 2)^4^. In addition, TIN2 recruits TPP1 (encoded by the gene ACD) that forms a heterodimer with POT1 (Protection of Telomeres 1)^13^. From the structural point of view, TIN2 is the central hub of the shelterin complex that links TRF1 and TRF2 homodimers with TPP1-POT1 heterodimer. Interaction domains of TIN2 that take part in TRF1, TRF2 and TPP1 binding are shown in Figure 1a.

**Figure 1.**
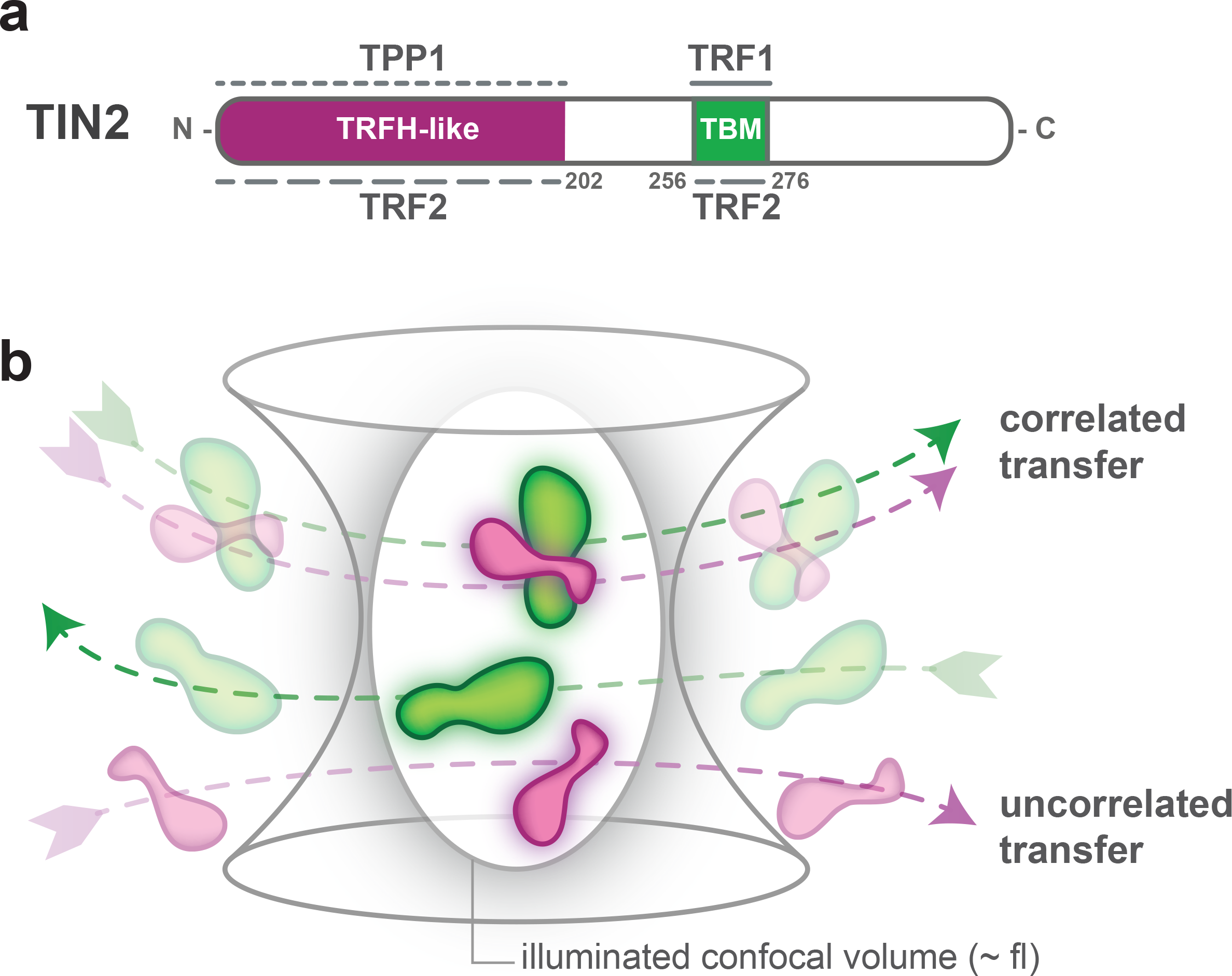
Interaction domains of TIN2 that interconnect TRF1, TRF2 and TPP1. (**a**) TRFH-like dimerization domain mediates interaction with TRF2 and TPP1; TBM - TRFH of TRF1/TRF2 binding motif. The more solid the line between TIN2 and the interacting proteins, the higher the mutual affinity^28,29^ (**b**) Scheme of how fluorescence cross-correlation spectroscopy detects bound proteins. When fluorescently labeled TIN2 and differently labeled TRF1 or TRF2 diffuse through the illuminated confocal volume, fluorescence signals fluctuations are recorded simultaneously. If TIN2 binds TRF1 or TRF2, the proteins move together, so they produce fluorescence intensity fluctuations of similar patterns for both fluorescence labels and the cross-correlation amplitude increases accordingly.

Regarding biological functions, TIN2 is essential for telomere length regulation mediated by TRF1^14,15^. TIN2 is required for TRF2-induced protection against ATM signaling pathway^16^ and POT1-meditated protection against ATR signaling pathway^17,18^. TIN2 deletion compromises the stability of both TRF1 and TRF2 at telomeres in cells^12,19^. TIN2 bridges TRF1 and TRF2 with TPP1 that recruits telomerase to telomeres^5,20^ and enhances telomerase processivity upon complexation with POT1^21–23^. The assembly of shelterin subunits around TIN2 is critical for the formation of structurally and biologically functional shelterin complex. Mutations in the gene of TIN2 have been implicated in approximately 15% of all known cases of *dyskarotis congenita* – a disease that results in defective telomere maintenance of adults^24,25^. Thus, the shelterin assembly mechanism and the stoichiometry of associated proteins are in the center of interest of telomere research community.

The overall shelterin stoichiometry on telomeres is known *in vivo*^26^. Newly, the *in vitro* stoichiometry of an assembled core complex comprising TRF2, TIN2, TPP1, and POT1 was revealed to be 2:1:1:1, respectively^27^. Previous studies described the structure and binding affinity of peptides representing interaction regions that take part in TRF1 and TRF2 binding to TIN2^28^. Very recently, the structure of the isolated interacting domains of TRF2, TIN2 and TPP1 has been determined^29^.

On the contrary, very little is known about how full-length TRF1 and TPP1 affect other full-length shelterin proteins during their assembly. The quantitative studies of shelterin proteins moving freely in solution represent an experimental challenge connected with the comparable size of TRF1 and TRF2 that excludes using simple fluorescence polarization measurements. Recently more extensively introduced single-molecule approaches are powerful tools of assessing the functional states of a molecular system as has been demonstrated by assessing DNA-repair complex assembly and dynamics^30,31^.

To reveal the structure, assembly and function of telomeric proteins and telomerase, single-molecule approaches could be applied as carefully reviewed by Parks and Stone^32^. However, classical TIRF (Total Internal Reflection Fluorescence) microscopy based single-molecule approaches are often limited to the area near the surface, as studied molecules are attached to the surface. On the other hand, if we use confocal scanning microscopy, we can measure interactions of fluorescently labeled proteins moving freely in solution regardless of the distance from the surface.

In this study, we took advantage of Fluorescence Cross-Correlation Spectroscopy (FCCS) - a single-molecule method that is based on an evaluation of the interdependence of time-resolved fluctuations of two different fluorophores by confocal microscopy^33^. FCCS monitors simultaneous fluorescence signals of two differently labeled proteins that diffuse through the confocal volume of a microscope objective (Figure 1b)^34,35^. FCCS has previously been extensively used to describe assembly of oligomeric calcium/CaM-dependent kinase II and calmodulin by the Schwille laboratory^33^.

We used FCCS to monitor protein binding based on the change in relative cross-correlation of differently labeled TIN2, TRF1 and TRF2 *in vitro* and to address the following hypotheses.

First, we wanted to know whether both TRF1 and TRF2 bind TIN2 simultaneously or if there is an order preference during the shelterin subcomplex assembly. Additionally, we tested the hypothesis that TPP1 binding to TIN2 may improve TRF2-TIN2 interaction and could enable TIN2 to interact simultaneously with TRF1 and TRF2 as has been suggested by the Songyang laboratory^36^. Finally, we wondered if we could suggest an interaction model of TRF1, TRF2, TIN2, and TPP1 assembly and correlate the model with available structural data and biological functions of shelterin proteins.

We found that TRF1 induces TRF2 release from TIN2. We also described that TPP1, upon binding to TIN2, improves TIN2’s binding capacity so the complex TIN2-TPP1 can accommodate both TRF1 and TRF2. Thus, we confirmed, at the single-molecule level, that TPP1 is essential for the formation of the stable TRF1-TIN2-TPP1-TRF2 complex. We suggest a mechanism that explains the exclusivity of TRF1-TIN2 interaction along with the requirement of TPP1 for simultaneous binding of TRF1 and TRF2 to TIN2. This work is, to our knowledge, the first single-molecule study describing full-length proteins TRF1, TRF2 and TIN2 during shelterin assembly in solution. The application of new experimental approaches enabled us to describe how the arrangement and structure of functional subcomplexes contribute to the role of complete shelterin in telomere protection.

## Results

### TRF1 replaces TRF2 bound to TIN2

We wanted to know whether TIN2 can accommodate both TRF1 and TRF2 simultaneously. According to previous measurements of Hu et al. and Chen et al.^28,29^ along with our MST assays (Figure 2-figure supplement 2) we allowed to form stable complexes of TRF1-TIN2 and TRF2-TIN2 at micromolar concentrations. We prepared protein complexes in 2:1 stoichiometry for TRF1 or TRF2 and TIN2 according to Lim et al.^27^, in all experiments within this study. TRF1 or TRF2 (20 nM) labeled with red fluorophore Alexa Fluor 594 in complex with TIN2 (10nM) labeled with the green fluorophore Alexa Fluor 488 were used in our FCCS measurements.

In the first experiment, we titrated dual-labeled TRF2-TIN2 complex with unlabeled TRF1 to the final concentration 80 nM (Figure 2a).

**Figure 2.**
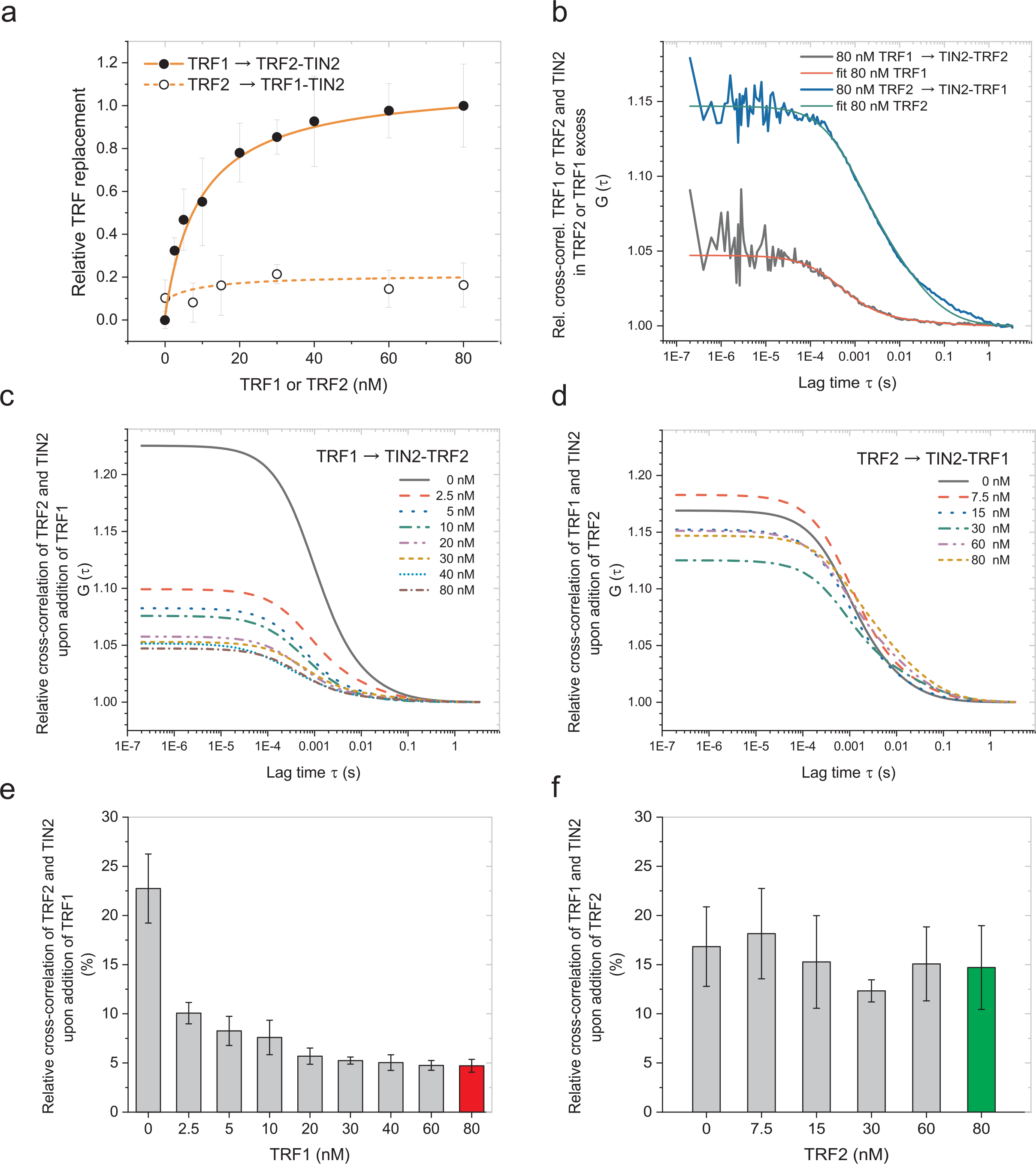
Relative cross-correlation decrease indicates that TRF1 releases TRF2 from TIN2. (**a**) The relative cross-correlation of fluorescently labeled TRF2 (20 nM) and TIN2 (10 nM) or TRF1 (20 nM) and TIN2 (10 nM) was measured upon addition of unlabeled TRF1 or TRF2 (0-80 nM), respectively. The proportional amplitude change (decrease) is presented as a “binding” curve. TRF1 addition disrupts TRF2-TIN2 (2:1) relative cross-correlation, but TRF2 addition to the preformed TRF1-TIN2 (2:1) complex does not affect relative cross-correlation (it stays constant). (**b**) Raw relative cross-correlation curves and their fits for TRF1 or TRF2 addition at the highest used concentration (80 nM) to TRF1-TIN2 or TRF2-TIN2, respectively. (**c**) Fits of relative cross-correlation curves upon addition of TRF1 show gradual decrease in amplitude of TRF2-TIN2 relative cross-correlation in accordance with TRF1 concentration increase. (**d**) Fits of relative cross-correlation curves upon addition of TRF2 to TRF1-TIN2 demonstrate that the amplitude is unchanged in the whole concentration range. (**e**) TRF2-TIN2 relative cross-correlation amplitudes decrease upon addition of TRF1. (**f**) Upon TRF2 addition, the relative TRF1-TIN2 cross-correlation amplitude stays intact and oscillates within the standard error.

We observed that TRF1 decreased the relative cross-correlation between TIN2 and TRF2 immediately after TRF1 addition to total 2.5 nM concentration. The decrease of relative cross-correlation suggested that TRF1 replaced TRF2 in complex with TIN2 (Figure 2a, c, e). Overall, our results showed that TRF1 addition diminished the relative cross-correlation of TRF2-TIN2 to the level of negative control (Figure 2b, c, e, Figure 2-figure supplement 1). Thus, the decrease of relative cross-correlation between TRF2 and TIN2 in the presence of TRF1 demonstrated that TRF1 released TRF2 from TIN2.

### TRF2 showed no influence on TRF1-TIN2 complex

In the next sets of experiments, we used a reverse arrangement where TRF1 labeled with red fluorophore (Alexa Fluor 594) was allowed to bind TIN2 labeled with green fluorophore (Alexa Fluor 488). Subsequently, we added unlabeled TRF2 gradually and monitored if TRF2 can disturb the complex TRF1-TIN2. We measured relative cross-correlation of labeled TRF1 and TIN2 at each concentration of unlabeled TRF2 (Figure 2d). The relative cross-correlation of TRF1 and TIN2 remained high and stable at TRF2 concentration up to 80 nM (Figure 2a, b, d and f). In other words, when we added TRF2 to the preformed complex TRF1-TIN2, we detected no significant decrease of relative cross-correlation between TRF1 and TIN2. The minimal effect of TRF2 presence on TRF1-TIN2 relative cross-correlation suggested that TRF2 did not disturb the TRF1-TIN2 complex.

### TRF2 has no effect on DNA binding affinity of TRF1-TIN2

We wanted to know whether the presence of full-length TRF2 affects TRF1 binding to telomeric DNA when TRF1 is in complex with TIN2. To analyze how mutual TRF1, TRF2 and TIN2 interactions affect DNA binding affinity, we employed fluorescence anisotropy (Figure 3). For DNA binding studies describing affinity to telomeric duplex R5 comprising five telomeric repeats (for exact sequence of R5 see Materials and Methods), we used stoichiometric combination of TRF1, TRF2 and TIN2, 2:2:1, respectively, as has been suggested previously^26,27^. R5 should feasibly accommodate both TRF1 and TRF2 simultaneously. Our quantitative binding data revealed that DNA binding affinity of the stoichiometric combination of TRF1, TRF2 and TIN2 is similar to DNA binding affinity of the combination TRF1 and TIN2. In other words, TRF2 did not affect DNA binding affinity of TRF1-TIN2.

**Figure 3.**
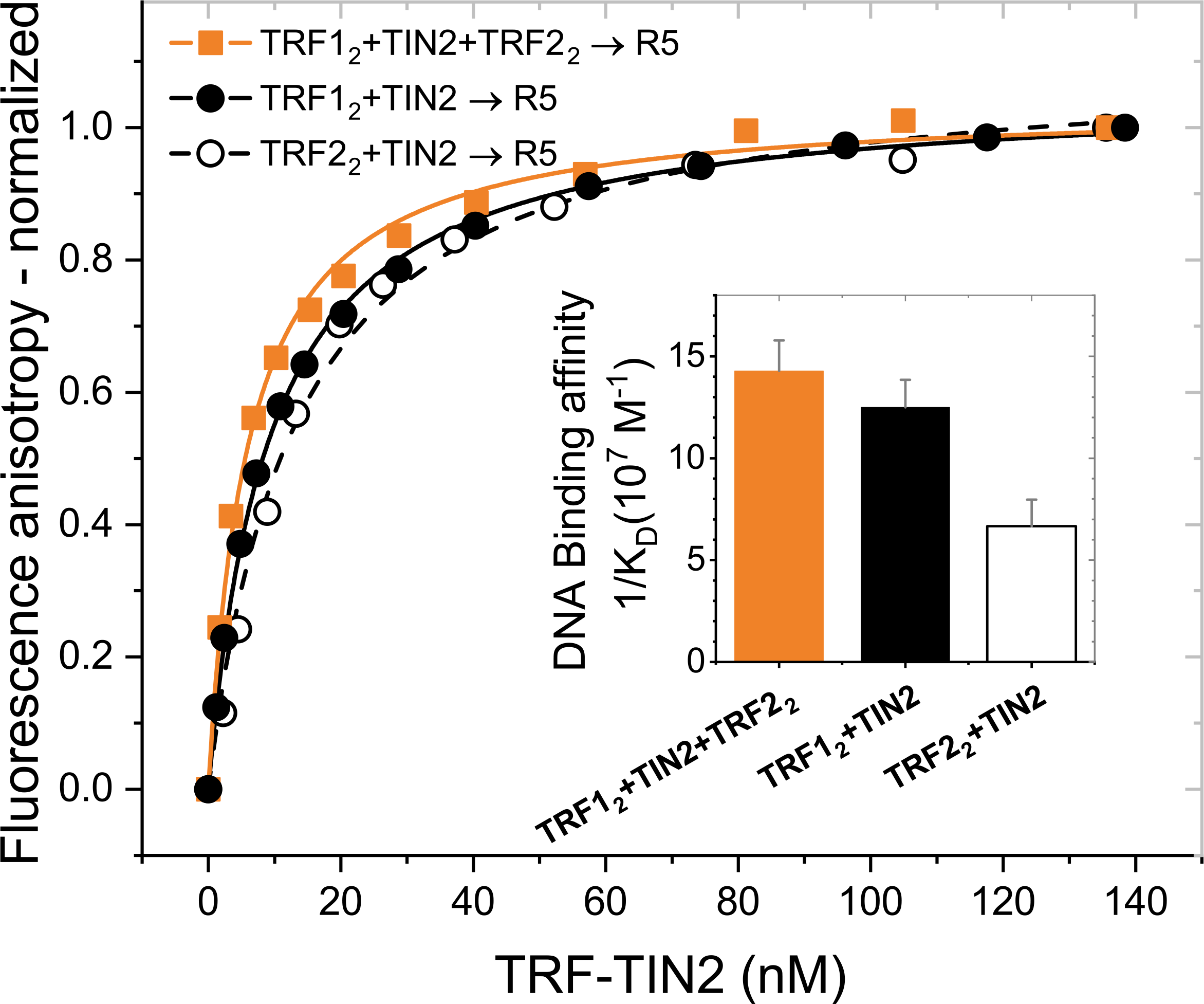
DNA binding affinity of stoichiometric combination of TRF1:TIN2:TRF2 (2:1:2) is similar to DNA binding affinity of stoichiometric combination of TRF1:TIN2 (2:1). TRF1 (5 μM) and/or TRF2 (5 μM) were incubated with TIN2 (2.5 μM) in 50 mM sodium phosphate, pH 7.0, 50 mM NaCl at 25°C. Protein solutions were titrated to Alexa Fluor 488 labeled DNA duplex (7.5 nM) in 50 mM sodium phosphate, pH 7.0, 50 mM NaCl at 25°C. Binding to telomeric DNA duplex R5 containing five telomeric repeats was measured by fluorescence anisotropy. The presented biding isotherms are averages of at least five independent experiments with standard deviation lower than 3% for each presented data point. The binding curves were normalized for the comprehensive comparison. Data were fit to a one-site binding model by non-linear regression to obtain equilibrium dissociation constants (K_D_). The inset bar plot shows reciprocal K_D_ values corresponding to DNA binding affinity.

### TRF2 and telomeric DNA create aggregates that prevent single-molecule assessment

Further, we wanted to assess by FCCS the TRF1-TIN2-TRF2 interactions in the presence of DNA. Unfortunately, we observed that TRF2 and telomeric DNA duplex R5 formed aggregates that forbade effective single-molecule monitoring of TRF2 interactions (Figure 4, Figure 4-figure supplement 1). The aggregates became clearly visible by fluorescence microscopy from TRF2 concentration 100 nM. We have obtained the same result when we carried out the measurements with TRF2 lacking the basic B-domain.

**Figure 4.**
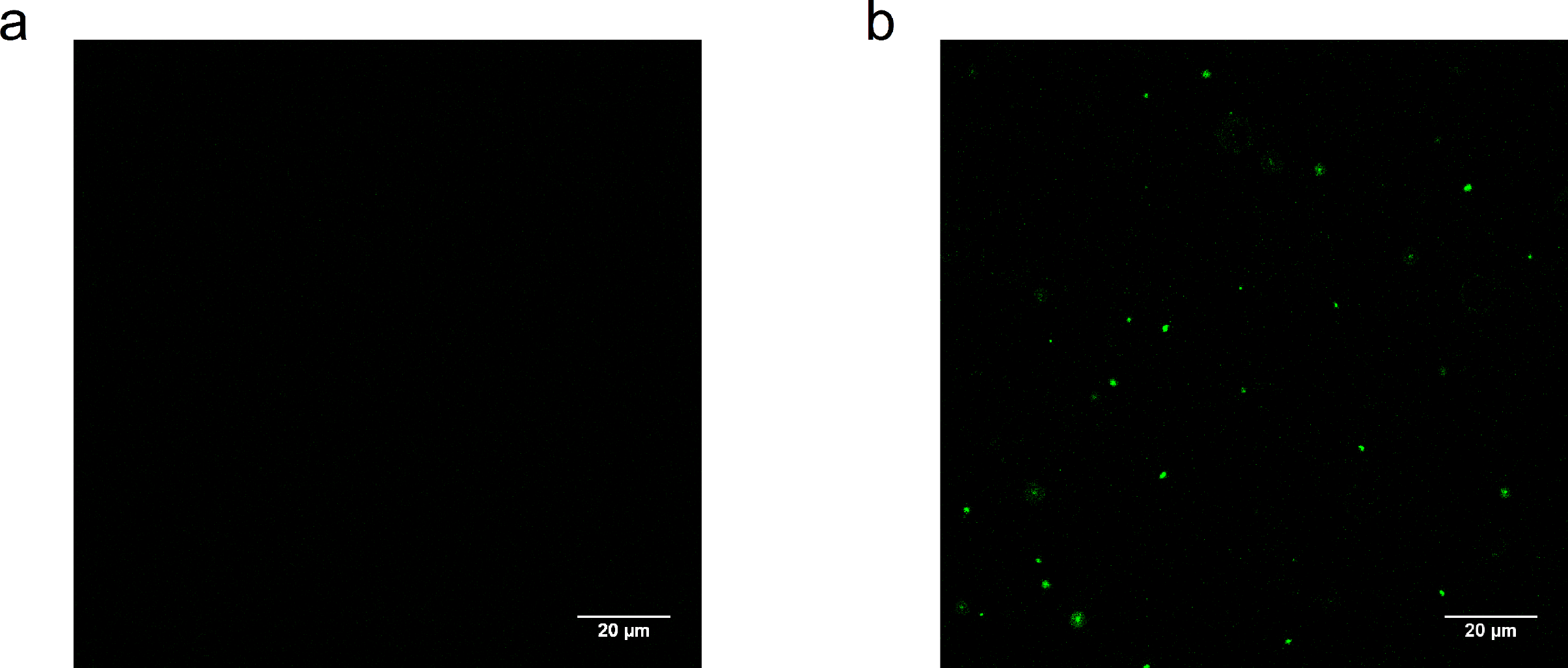
Upon binding to human telomeric DNA R5, TRF2 forms aggregates visible by fluorescence microscopy. (**a**) Fluorescently labeled DNA containing five telomeric repeats (R5) was diluted in 50 mM sodium phosphate buffer, pH 7.0 with 50 mM NaCl to the final concentration of 10 nM. Without TRF2 no visible signal was detected. (**b**) After TRF2 addition to 400 nM, fluorescence signals of aggregations were clearly visible.

### TPP1 enables TIN2 to accommodate both TRF2 and TRF1 simultaneously

We wondered whether another human telomeric protein TPP1 improves the stability of TRF1-TIN2-TRF2 complex consisting of full-length proteins. Hu et al. suggested that the C-terminal domain of TPP1 is responsible for its binding to TIN2 and that TPP1 stabilizes TIN2-TRF2 interaction^29^. Recombinant human TPP1 with an N-terminal deletion, TPP1 (89-554), was used, as full-length TPP1 purity was insufficient for single-molecule experiments. TPP1 (89-554) was chosen because the 88 N-terminal residues of TPP1 are functionally dispensable in human cells and are not conserved among TPP1 proteins of different organisms^23,37,38^. Additionally, TPP1 (89-554) still contains N-terminus of OB domain that is critical for telomerase activity^39^. For simplicity, we hereafter use TPP1 to represent TPP1 (89-554) unless stated otherwise. We incubated fluorescently labeled TRF2 and TIN2 along with unlabeled TPP1 at room temperature. Then, we added unlabeled TRF1 to see whether TRF1 can still replace TRF2 from the TIN2-TPP1-TRF2 complex. As Figure 5 shows, relative cross-correlation between TRF2 and TIN2 remained at high levels for all concentrations of TRF1. The unchanged relative cross-correlation suggested that TRF2 remains bound to TIN2 in TRF1 and TPP1 presence. The formation of the complex TRF1-TIN2-TPP1-TRF2 was confirmed by a complementary FCCS experiment where TRF1 and TRF2 were labeled by Alexa Fluor 488 and Alexa Fluor 594, respectively. First, unlabeled TIN2 was added to the mixture of labeled TRF1 and TRF2. After TIN2 addition, we observed no significant change in amplitude of relative cross-correlation between TRF1 and TRF2 (Figure 5-figure supplement 1). On the contrary, when we added preformed complex of TIN2-TPP1 into TRF1 and TRF2 mixture, we detected substantial increase of relative cross-correlation between TRF1 and TRF2 (Figure 5-figure supplement 1).

**Figure 5.**
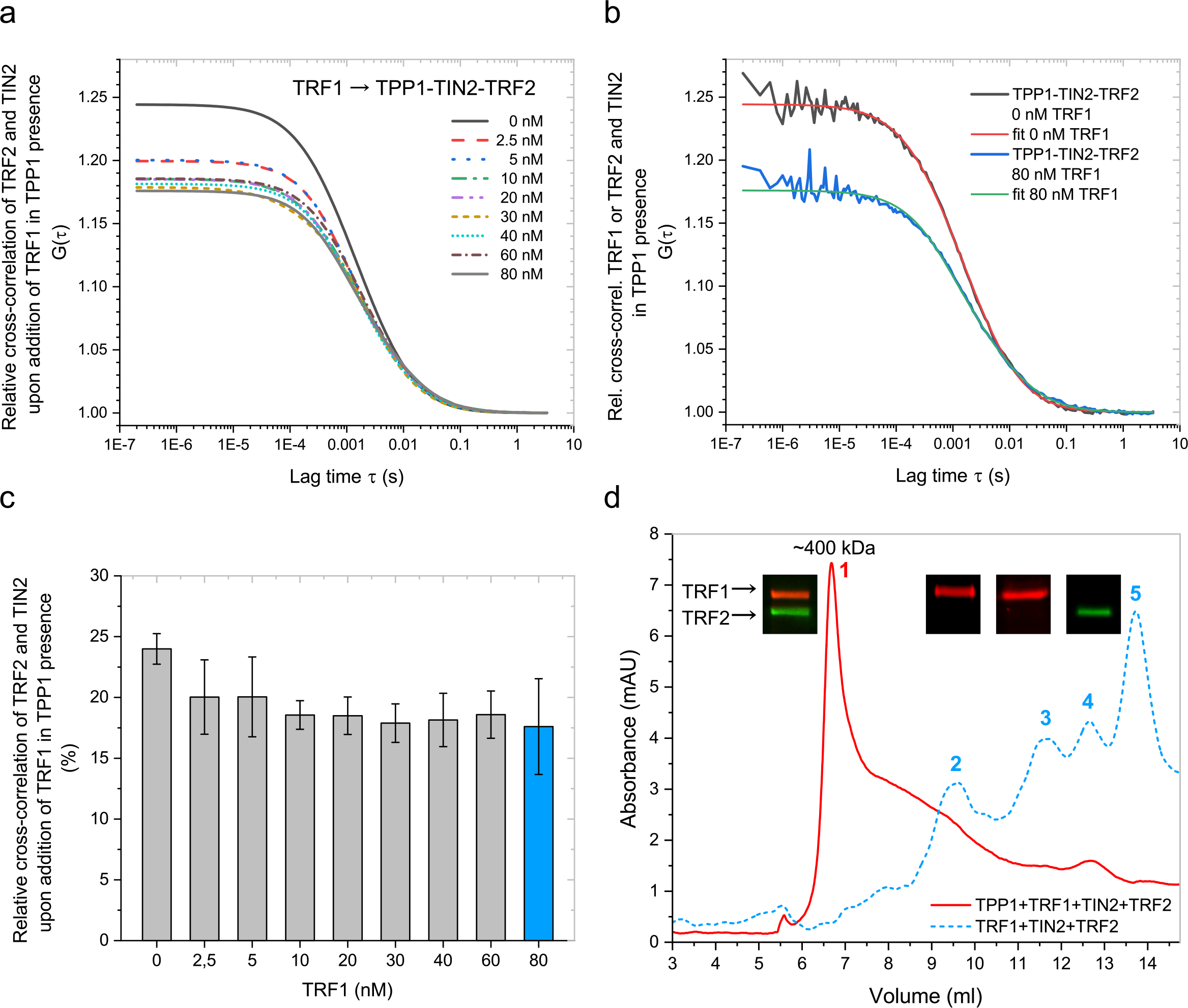
TRF2 remained bound to TIN2-TPP1 complex in TRF1 presence. (**a**) Alexa Fluor 488 labeled TIN2 (1 μM) and Alexa Fluor 594 labeled TRF2 (2 μM) were incubated in stoichiometric ratio with unlabeled TPP1 (1 μM) at 25°C to form a complex. The sample was diluted to the final complex concentration 20 nM. Fits of relative cross-correlation curves upon addition of TRF1 to TRF2-TIN2-TPP1 complex show high amplitudes regardless of presence of TRF1 up to 80 nM concentration. (**b**) Representative raw cross-correlation curves and fits for TRF2 and TIN2 in TRF2-TIN2-TPP1 complex recorded in the absence of TRF1 or in the presence of TRF1 at 80 nM concentration from (A). (**c**) In the presence of TPP1, TRF2-TIN2 relative cross-correlation amplitude remains high upon addition of TRF1, indicating that TPP1 stabilizes TRF2-TIN2 complex and allows TRF1-TIN2-TRF2 complex formation. (**d**) Representative size-exclusion chromatography traces of TRF1, TIN2, TPP1, TRF2 (solid line) and TRF1, TIN2, TRF2 (dashed line). Proteins were mixed and incubated in the same order as they appear. Superdex™ 10/300 GL column with 50mM NaCl, 50mM phosphate pH 7.0 as mobile phase was used for the chromatographic separation. TRF1 and TRF2 were labeled by AlexaFluor 594 and AlexaFluor 488, respectively. Fractions corresponding to numbered peaks were collected and analyzed on SDS-PAGE. Labeled proteins in gels were detected using Typhoon™ FLA 9500 detection system. Peak 1 contains both labeled proteins within TRF1-TIN2-TPP1-TRF2 complex. Peak 2 represents TRF1-TIN2, peak 3 - TRF1, peak 4 - TRF2, peak 5 - TIN2.

When we mixed TRF1, TIN2, TRF2 and TPP1 together, we observed comparable relative cross-correlation between TRF1 and TRF2 as the relative cross-correlation between TRF2 and TIN2 in previous experimental setup (compare Figure 5 and Figure 5-figure supplement 1). The high relative cross-correlation level in both experimental arrangements verified that TRF1-TIN2-TPP1-TRF2 complex was formed.

To confirm that proteins form a stable complex, we recorded size-exclusion chromatography profiles of mixture comprising TRF1, TIN2 and TRF2 with or without TPP1 (Figure 5d). Only in presence of TPP1 we observed a high-molecular peak that corresponds to the assembled protein complex (compare the solid red line and dashed blue line in Figure 5d). We analyzed collected chromatographic fractions by SDS gel electrophoresis. As TRF1 and TRF2 were labeled by different fluorophores, we identified electrophoretic bands corresponding to TRF1 and TRF2 in fluorescence densitograms (insets in Figure 5d). The fluorescence intensity profiles showed that TRF1 and TRF2 formed a complex only if TPP1 and TIN2 were present.

Additionally, we have carried out control measurements with TIN2 mutants that were unable to bind TPP1 or TRF2 on the N––terminal part of TIN2. We have prepared two TIN2 point mutants - A15R and A110R. A15R mutation of TIN2 prevents TPP1 binding; A110R mutation of TIN2 prevents TRF2 binding to N-terminal binding site of TIN2, as revealed by Hu et al.^29^. FCCS studies showed that both mutations of TIN2 restricted the assembly of shelterin core complex (Figure 5-figure supplement 2). We found that A110R TIN2 was unable to form complex with TRF2 even in TPP1 and TRF1 presence (Figure 5-figure supplement 2a). The inability of A110R TIN2 to bind TRF2 indicates that N-terminal-binding site of TIN2 is critical for stable accommodation of TRF2 into the complex. FCCS measurements with the second mutant revealed that A15R TIN2 with impaired TPP1 binding ability was unable to cross-correlate with TRF2 in the presence of TRF1 and TPP1 (Figure 5-figure supplement 2b). Taken together, FCCS experiments with TIN2 mutants supported the view that the N-terminal binding domain of TIN2 is essential for the cooperative binding of TRF2 and TPP1 to TIN2.

In summary, our combined single-molecule and ensemble analyses of full-length shelterin core proteins supported the view that TPP1 enables TIN2 to bind both TRF1 and TRF2 together and form stable TRF1-TIN2-TPP1-TRF2 complex.

## Discussion

Our findings of how human telomeric proteins TRF1 and TPP1 affect the formation of core shelterin complex TRF1-TIN2-TRF2 have provided new insights into the assembly of full shelterin complex at the single-molecule level. In addition, our study has contributed to address the following biological questions about human shelterin:

(i) What are the arrangements of shelterin proteins TRF1, TRF2 and TIN2 in solution? (ii) How does the shelterin core TRF1-TIN2-TRF2 assemble? (iii) How can TPP1 affect TRF1-TIN2-TRF2 complex formation? We address these questions below.

Our quantitative biophysical observations clearly showed that TIN2 bind either TRF1 or TRF2. Thus, two independent complexes TRF1-TIN2 and TRF2-TIN2 appear in solution. The observation of exclusive binding of TRF1 or TRF2 to TIN2 is in agreement with previous studies. O’Connor et al. have suggested that TRF1-TIN2 and TRF2-TIN2 occur as separate sub-complexes based on immunoprecipitation studies^36^. Additionally, when we take into consideration available information about the positions of interacting domains, domain structure and quantitative binding characterizations, we can rationalize why TRF1 replaces TRF2 on TIN2. So far, it seems that shelterin proteins form hetero-multimer complexes through a selective domain-domain-interaction mechanism.

As Chen et al. showed, both TRF2 and TRF1 can bind one common binding site TBM (TRFH-binding motif, Figure 1a) on TIN2^28^. TBM at the C-terminus of TIN2 is a well-structured 256-276 region that interacts with TRFH domain of both TRF1 and TRF2. The surface of TBM matches better the hydrophobic interface of TRFH domain of TRF1 than the polar interface of TRFH domain of TRF2. The different structural arrangements of interaction interfaces prompt that TRFH domain of TRF1 binds a peptide representing the TBM region of TIN2 with higher binding affinity than the TRFH domain of TRF2^28^.

In addition, there is another well-structured binding site at TIN2’s N-terminus (TRFH-like) where TRF2 binds with higher affinity compared to the previously mentioned common TRF1/TRF2 binding site TBM. If we consider that interactions between proteins occur mainly through the minimal identified domains, we may expect similar binding affinity for full-length proteins. Thus, TRF1 may form a complex with TIN2 more readily than TRF2. The higher binding affinity of TRF1 to TIN2 causes higher preference for the formation of complex TRF1-TIN2 compared to TRF2-TIN2. The ensemble binding affinity of full-length TRF1 to TIN2 with K_D_ 240 nM and TRF2 to TIN2 with K_D_ 360 nM have been measured by Microscale thermophoresis (Figure 2-figure supplement 2). Here obtained higher binding affinities for full-length proteins than affinities for isolated domains measured by Chen et al.^28^ and Hu et al.^29^ might suggest that additional hydrophobic and hydration effects promote full-length protein interactions^40^.

If we consider that TRF1 and TRF2 concentrations are similar *in vivo*^26^, we suppose that TRF1-TIN2 complex incidence should prevail over TRF2-TIN2 complex occurrence significantly. Additionally, if there was no other binding site on TIN2 for TRF2 or a binding regulation mechanism, the probability of forming complex TRF1-TIN2-TRF2 would be negligible. The second binding site for TRF2 on TIN2 (TRFH-like domain, Figure 1a) should allow tri-functional complex TRF1-TIN2-TRF2 formation.

In this context, our finding that TRF1 can substitute TRF2 when bound to TIN2 was rather unexpected at first glance (Figure 2a, c, e). However, we can rationalize TRF2 displacement when we consider that TRF1 binding affinity to TIN2 is ten-fold higher^29^ than TRF2 binding to TBM on TIN2^28^, as supported by our MST measurements with full-length proteins (Figure 2-figure supplement 2). Let us assume that TRF2 binds TRFH-like domain of TIN2 in TRF1 absence, as TRF2 should occupy the binding site with the highest affinity at first. When TRF1 appears in TIN2-TRF2 complex proximity, TRF1 binds TBM site on TIN2.

Why is TRF2 released upon TRF1 binding when there are two independent binding sites for TRF1 and TRF2? One straightforward explanation would be that TRF1 bound to TIN2 presents a steric hindrance that disturbs optimal interaction surface between TRF2 and TIN2. The second explanation could be that TRF1 induces allosteric changes in TRFH-like domain of TIN2 that disable TRF2 binding to TIN2. Moreover, there could be a combination of both – structural restrictions and allosteric changes. Nevertheless, TRF2 binding to TRF1-TIN2 has to be promoted to form the stable core complex TRF1-TIN2-TRF2.

As the first, O’Connor et al. suggested that TPP1 promotes the interaction between TIN2 and TRF2^36^. Recently, Hu et al. have shown that the binding of TPP1 interacting domain to TIN2’s TRFH-like domain allosterically changes TRF2 binding site on TRFH-like domain of TIN2^29^. Furthermore, Hu et al. revealed that the binding affinity between minimal interaction domains of TPP1-TIN2 and TRF2 was increased almost three-fold if compared to the interaction without TPP1^29^.

As TPP1 induces allosteric changes on TIN2 that enable TRF2 binding and increase TRF2 binding affinity, TPP1 acts as an assembly activator that is required for the stable formation of TRF1-TIN2-TPP1-TRF2 complex.

Our single-molecule FCCS results support the view that TPP1 acts as a shelterin assembly activator. We corroborated that TRF2 remained bound to TIN2 in TRF1 presence and TRF1-TIN2-TRF2 complex was formed when TPP1 was bound to TIN2 (Figure 5a, Figure 5-figure supplement 1). We summarize our recent results within the framework of the present state of knowledge of shelterin core assembly in the model below.

### Model of core shelterin assembly

We propose a model of how shelterin core proteins assemble in solution. The model takes into consideration that TRF1 and TRF2 form homodimers^41,42^, as the homodimerization exclusivity of TRF1 and TRF2 is a functional requirement that facilitates separation of different functions for the both TRF proteins of similar domain structure. The model also reflects the stoichiometry of shelterin proteins that has been revealed by the de Lange and the Cech laboratories^26,27^.

The recommended model suggests that if TRF1 occupies the TBM binding site on TIN2, TRF2 binding to TIN2 is compromised and only TRF1 remains bound to TIN2 (Figure 6). When TPP1 binds TIN2, TIN2 is allosterically changed and its N-terminal binding site becomes active. Then, TRF2 can bind TIN2 also in TRF1 presence. Thus, TPP1 activates the N-terminal binding site for TRF2 on TIN2 and enables TIN2 to accommodate both TRF1 and TRF2 simultaneously. The model suggests that the protein order during shelterin core self-assembly in solution is TRF1 > TIN2 > TPP1 > TRF2.

**Figure 6.**
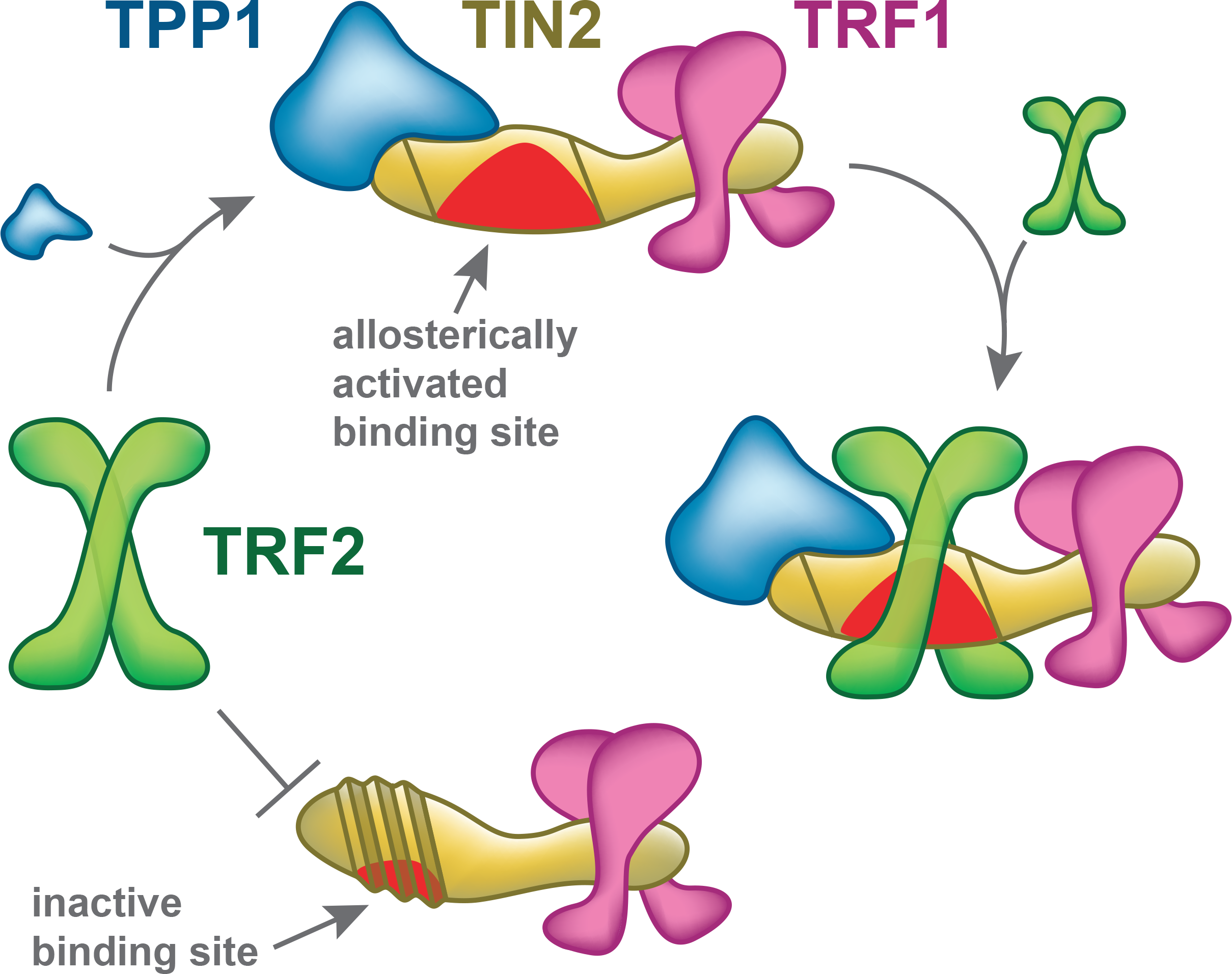
A model for sequential assembly of shelterin core complex TRF1-TIN2-TRF2. TRF1 prevents TRF2 from binding to TIN2 if TPP1 is absent, as TRF1 occupies the preferential binding site on TIN2 (lower part of the scheme). On the contrary, when TPP1 binds TIN2, TPP1 induces allosteric structural changes that open the second binding site on TIN2 and the binding site for TRF2 becomes active (upper part of the scheme). TRF2 binds TIN2-TPP1 along with TRF1 (middle part of the scheme). A stable shelterin core complex TRF1TIN2TPP1TRF2 is formed.

Additionally, the model explains the unique ability of TRF1 to exclude TRF2 from the complex TRF2-TIN2. Our model suggests that TRF1-TIN2 is an initial complex based on the highest affinity between TIN2 and TRF1 among shelterin proteins^28^. Moreover, TRF1-TIN2 preferential binding explains why it is not possible to prepare TRF1-TIN2-TRF2 complex without TPP1. TPP1 must induce allosteric structural changes on TIN2 to open N-terminal TRF2 binding site^29^.

The induced structural changes allow tighter binding of TRF2 to TIN2 without the compromising effect of the prebound TRF1.

The proposed model suggests that TRF1 and TRF2 dimers are bound to TIN2 monomer. The equimolar TRF1:TRF2 ratio is in accordance with our size-exclusion chromatography analysis that showed the same fluorescence intensity of labeled TRF1 and labeled TRF2 in complex with TIN2 and TPP1 (Figure 5d, red solid line). The equimolar ratio TRF1:TRF2 along with previous observation of the Cech laboratory that TRF2 forms complex with TIN2 in 2:1 stoichiometry^27^ suggest that TRF1:TIN2 complex has 2:1 stoichiometry.

The model with two TRF2 binding sites on TIN2 has been supported by our FCCS experiments with mutated variants of TIN2 (Figure 5-figure supplement 2). We observed no cross-correlation for TIN2 mutants that were unable to bind TPP1 or TRF2 on N-terminus of TIN2. The point mutations of TIN2 prevented the complexation of TRF1 and TRF2. The model supports the view that the N-terminal binding domain of TIN2 is essential for the cooperative binding of TRF2 and TPP1 to TIN2 and promoting assembly of TRF1-TIN2-TRF2-TPP1 complex.

The proposed model is applicable also to mechanisms where proteins first form weak transient complexes, and then depend on the additive energies of binding and allosteric changes provided by partner proteins to generate higher specificity. Finally, the intrinsic dynamics of TRF1 and TRF2 could be important for regulating the assembly and disassembly of shelterin complexes and exchanging between capped and uncapped telomere structures^43^.

The question is how the suggested model of protein assembly corresponds to the situation when DNA appears in solution. We speculate that telomeric DNA could serve as a scaffold to which TRF1 and TRF2 bind with high affinity^11,44^. Thus, DNA may promote the assembly of TIN2, TRF1 and TRF2 without TPP1. In cellular environment, the situation could be more intricate if telomeric DNA is folded to quadruplexes that might prevent binding TRF1 and TRF2. If DNA is in quadruplex form, POT1 has to be engaged to unfold quadruplexes^45^ and allow formation of double-stranded regions and single-stranded junctions on telomeric DNA – TRF1 and TRF2 primary binding sites. As POT1 dimerizes with TPP1 readily^29,46^ and TPP1 contributes to higher affinity of POT1 to DNA^23^, one could anticipate that TPP1 with POT1 might be attached to a flexible single-stranded telomeric overhang. TPP1 would be then available to mediate assembly of TIN2 and TRF1 together with TRF2 bound on relatively rigid double-stranded DNA. The flexibility of single-stranded DNA would allow interaction of POT1/TPP1 with core shelterin TRF1-TIN2-TRF2 bound in close proximity. As the interactions between protein subunits and DNA likely occur simultaneously, in the strictest sense, initial mutual binding affinities of shelterin proteins may not necessarily describe the order of their assembly. Our results together with recent structural and functional studies advocate separation between initial binding and final locking interactions leading to stable but still dynamic shelterin assembly.

In summary, our study with full-length human telomeric proteins brings new information about assembly of shelterin subcomplexes in solution. Our results with the core shelterin complex TRF1-TIN2-TRF2 extend the knowledge, so far limited to previous structural and functional studies of shelterin assembly that have been carried out without TRF1. For the first time, we applied single-molecule approaches to monitor full-length TRF1, TRF2, TPP1 and TIN2 during their assembly in solution. The presented studies describe how the mutual arrangement of functional subcomplexes of telomeric proteins contribute to the role of the whole shelterin in telomere protection. The next challenging tasks will be to describe the effects of the DNA scaffold on shelterin assembly and monitor the dynamics of telomeric proteins associations in living cells.

## Material and Methods

### Cloning, expression and purification of TRF1, TRF2, TIN2 and TPP1

The cDNA sequences of TRF1, TRF2, TIN2, and TPP1 were synthesized by Source BioScience and cloned to pDONR/Zeo vector (Life Technologies) using two sets of primers and BP clonase enzyme mix from Gateway technology (Life Technologies). The resulting plasmids were cloned into different expression vectors in a recombination reaction using LR clonase enzyme mix (Life Technologies) and expressed as His-tagged proteins in different strains of Escherichia coli (pUbiKan_X105_TRF1 and pHGWA_TRF2 in BL21(DE3), pTriEx4_TIN2 in C41 and pRbXKan_x105_TPP1-FL (89. - 554.) in BL21(DE3) RIPL. BL21(DE3) and C41 cells harboring TRF1, TRF2 and TIN2 were grown in Luria-Bertani medium, BL21(DE3) RIPL cells with TPP1 were grown in Terrific Broth medium; containing 50 μg.ml^−1^ kanamycin (TRF1, TPP1) or 100 μg.ml^−1^ ampicillin (TRF2, TIN2) or 34 μg.ml^−1^ chloramphenicol (TPP1) at 37°C until A_600_ reached 0.5 (TPP1) or 1.0 (TRF1, TRF2, TIN2). The cells were cultured for 3 h at 15°C (TRF1, TPP1) or 25°C (TRF2, TIN2) after the addition of IPTG to the final concentration of 0.5 mM (TRF1, TIN2, TPP1) or 1 mM (TRF2). Cells were collected by centrifugation (8 000 g, 8 min, 4°C).

The pellet was dissolved in lysis buffer containing 50 mM sodium phosphate, pH 8.0, 500 mM NaCl, 10 mM imidazole (TRF1, TRF2, TIN2) or 20 mM imidazole (TPP1), 0.5% Tween-20 (TRF1, TRF2, TIN2) or 0.5% Triton X-100 (TPP1), 10% glycerol, protease inhibitor cocktail cOmplete tablets EDTA-free (Roche). The cell suspension was sonicated for 3 min of process time with 1 s pulse and 2 s of cooling on ice (Misonix). Cell lysate supernatant was collected after centrifugation at 20 000 g, 4°C for 1 h. Proteins were purified by immobilized-metal affinity chromatography using TALON^®^ metal affinity resin (Clontech), where filtered supernatant (0.45 μm filter) was mixed with TALON^®^ beads and incubated 30 minutes. The proteins of our interest were eluted at 200 mM (TIN2), 300mM (TPP1) or 500 mM (TRF1, TRF2) imidazole in the same buffer without Tween-20 or Triton X-100. TRF2 and TIN2 were dialyzed into 50 mM sodium phosphate pH 7.0, 50 mM NaCl. TRF1 was loaded onto the HiLoad 16/600 column containing Superdex 200 pg (GE Healthcare Life Sciences) and resolved using 50 mM sodium phosphate buffer pH 7.0 with 200 mM NaCl. TPP1 expression tags were removed by HRV3-C protease at 4°C for 2 hours with 3 mM DTT. The final purification was on HiLoad Superdex 200 pg column (GE Healthcare Life Sciences) equilibrated with buffer containing 50 mM sodium phosphate pH 7.0 and 800 mM NaCl. The proteins were concentrated and the buffer was exchanged to 50 mM sodium phosphate pH 7.0, 50mM NaCl (TRF1, TRF2, TIN2) or 150mM NaCl (TPP1) by ultrafiltration (Amicon 3K/30K, Millipore).

The concentration of purified proteins was determined by the Bradford assay. We evaluated protein purity by SDS-polyacrylamide gels stained by Bio-Safe Coomassie G250 (Bio-Rad) (Figure 2-figure supplement 3). Western blotting and quantitative mass spectrometry analyses also confirmed the presence and purity of full-length proteins.

### DNA substrates

For the DNA binding affinity studies, DNA duplex R5 was prepared by annealing a fluorescently oligonucleotide (Alexa Fluor 488) with the sequence 5′ - GTTAGGGTTAGGGTTAGGGTTAGGGTTAGGGTTAG - 3′, respectively and its complementary strand. The sequence of R5 was designed in accordance to the optimal binding site of TRF2 defined by the de Lange laboratory^47^. The substrate was purified using a Mono Q 5/50 GL column (GE Healthcare) with a gradient of 50-1000 mM LiCl in 25 mM Tris-HCl, pH 7.5. All oligonucleotides were purchased from Sigma-Aldrich.

### Fluorescence anisotropy

Measurements of TRF1-TIN2 and TRF2-TIN2 binding to telomeric DNA duplex R5 labeled by Alexa Fluor 488 were performed on a FluoroMax-4 spectrofluorometer (Horiba Jobin Yvon, Edison, NJ). All experiments were carried out at 25°C. Fluorescence anisotropy was monitored at an excitation wavelength of 490 nm and emission wavelength of 520 nm. The slit width (both excitation and emission) for all measurements was 9 nm and the integration time was 1 s. The cuvette contained 1.4 ml of DNA duplex R5 (7.5 nM) in a buffer containing 50 mM sodium phosphate pH 7.0 and 50 mM NaCl. A protein mixture was titrated into the DNA solution in the cuvette and measured after a 2 min incubation. Fluorescence anisotropy at each titration step was measured three times and averaged with relative standard deviation always lower than 3%. The values of dissociation constants were determined by non-linear least square fits according to the equation r = r^MAX^ c/(*K*_d_+c) using ORIGIN^®^ 2018 (OriginLab, Northampton, MA) and confirmed by symbolic equation-based fitting using Dynafit^48^.

### Fluorescent protein labeling

Fluorescent protein labeling has been performed according to the protocol provided by the supplier with the following modifications. Alexa Fluor 488 or 594 labels (Molecular Probes - Invitrogen) in fourfold molar excess over protein were diluted in 1/10 volume of 1M sodium bicarbonate, fluorophores were mixed with protein (0.1 μg) and incubated for 1 hour at 4°C while stirring. The mixture was loaded on PD-10 columns (GE Healthcare) and eluted with 50 mM sodium phosphate pH 7, 50 mM NaCl. The protein degree of labeling higher than 85 % was confirmed by UV/Vis spectroscopy.

### Microscale thermophoresis (MST) assay

MST is an in-solution method for the quantitative description of molecular interactions. MST measurements were performed with a NanoTemper Monolith NT.115 instrument (NanoTemper Technologies). Fluorescently labeled TIN2 (Alexa Fluor 488) (10 μL) was incubated for 10 min on ice with different concentrations of TRF1 or TRF2 (10 μL), in 50 mM sodium phosphate buffer pH 7.0 with 50 mM NaCl. Then, 5 μL of the samples was loaded into premium treated capillaries, and MST measurements were collected at 12 °C at 30% blue-laser power and 60% light-emitting-diode power. The laser-on and laser-off intervals were 20 and 5s, respectively. MO.Affinity Analysis v2.2.4 software was used to fit the data and to determine apparent K_D_ values. All measurements were collected at least three times for two independently prepared sample sets.

### Theoretical Concept of FCS and FCCS

Fluorescence correlation spectroscopy (FCS) describes spontaneous fluorescence intensity fluctuations caused by rapidly diffusing molecules in a microscopic detection volume (about one femtoliter). FCS determines mobility and kinetics at single-molecule precision^33^. Fluorescence cross-correlation spectroscopy (FCCS) monitors two different fluorescence signals (two colors) collected at the same time and determines how their coincident fluctuations correlate to each other if the proteins are moving together. FCCS describes binding of measured proteins independently of diffusion rate. The cross-correlation function of two-color system, with one green-labeled particle *G* and the second with red-labeled particle *R* is described as follows

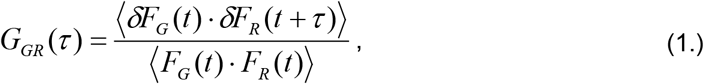

where *F_G_*(*t*) and *F_R_*(*t*) corresponds to fluorescence intensity fluctuations of individual signals (green and red) and τ represents lag time - the time period that two proteins stay in confocal volume together. A typical cross-correlation curve is a sigmoidal curve with the highest amplitude in time t(0). With longer lag time, the amplitude decreases, as the probability that two proteins are bound and stay in the confocal volume is lower with increasing time. The single-molecule nature of FCCS measurements causes that relative cross-correlation amplitudes of the same sample fluctuate within standard error, because we detect different number of molecules in confocal volume at different times. The cross-correlation fit with an appropriate model provides us with the degree of complexation of two fluorescently labeled molecules. Under ideal conditions (absence of FRET – Fluorescence Resonance Energy Transfer and spectral crosstalk of used fluorophores) the cross-correlation amplitude *G_GR_*(0) is directly proportional to the concentration of bound species^33,35^.

### Relative cross-correlation as a measure of binding

In FCCS experiments, we used TIN2 labeled with Alexa Fluor 488 and TRF1 or TRF2 labeled with Alexa Fluor 594, if not stated otherwise. For data evaluation, we refer to relative cross-correlation that was calculated as a ratio betwe en amplitude of cross-correlation function *G*_*TIN*2-*TRF*_(0) and auto-correlation function *G*_*TIN*2_(0). In other words, the relative cross-correlation corresponds to the proportion of cross-correlation amplitude related to the TIN2 auto-correlation amplitude. This approach also allowed us to normalize the data regarding their concentration-so we could directly compare the results of all measured FCCS experiments^33^.

### Conditions for Microscopy Imaging and Spectroscopy Measurements

All the FCCS measurements were performed with a confocal laser scanning microscope Zeiss LSM 780 using dedicated software ZEN Studio with additional FCS module. For excitation, the Ar^+^ laser was used with 488 and 561 nm continuous wave. The emission filters were tuned to minimize crosstalk between Alexa Fluor 488 and Alexa Fluor 594^49^. To determine the crosstalk, we carried out a FCCS measurement of a mixture of free fluorophores Alexa Fluor 488 and Alexa Fluor 594 that served also as a negative control with minimal cross-correlation (Figure 2-figure supplement 1). Confocal pinhole was fixed to 1 AU and it was additionally fine adjusted in x, y directions prior to FCCS measurement to maximize the count rate of fluorescence fluctuations. We used “FCS approved” water immersion objective (Zeiss 63x C-Apochromat NA 1.2 W Corr) which guarantees overlap of the point spread function (PSF) for different wavelengths and correction for any spherical aberration.

Fluorescently labeled TIN2 and TRF1 or TRF2 were incubated at 25°C at 1 μM concentration to form a complex. The sample was diluted to the final concentration that guaranteed the optimal amount of labeled proteins diffusing through the confocal volume at the same time (up to 5 proteins) corresponding to concentration 10-20 nM.

Prepared samples were incubated at room temperature, diluted with 50 mM sodium phosphate buffer pH 7.0 with 50 mM NaCl to the final volume of 200 μl and loaded into a μ-slide 8-well Glass Bottom Chamber (Ibidi). The sample covered the entire bottom of the well. Measurements were started immediately, without further incubation.

For each experiment, raw data containing 50 repetitions of 10-second acquisition were collected and averaged. This approach ensured that we collected enough data to obtain statistically significant values. The raw data were exported and analyzed with software QuickFit 3^50^. Correlation curves were averaged and fitted with two-component 3D Normal Diffusion model by solving the Levenberg-Marquardt nonlinear least-squares fitting routine. The data were visualized by ORIGIN^®^ 2018 (OriginLab, Northampton, MA). The experiments were performed in triplicate.

## Acknowledgements

The authors thank Petra Schwille and Thomas Weidemann for critical reviewing of FCCS data and their kind suggestions. The authors thank Blanka Pekarova for her assistance with size exclusion chromatography and Jitka Holkova for her help with microscale thermophoresis measurements. The authors are grateful to Victoria Marini for the critical reading of the manuscript. The Czech Science Foundation [16-20255S to C.H.] has supported this research. CIISB research infrastructure project LM2015043 funded by MEYS CR is gratefully acknowledged for the financial support of the measurements at the CF Proteomics and CF Biomolecular Interactions and Crystallization. The research has been carried out with institutional support of the Ministry of Education, Youth and Sports of the Czech Republic under the project CEITEC 2020 (LQ1601).

---

**Figure 2-figure supplement 1**. Relative cross-correlations of negative and positive control reactions. The negative control - free dyes Alexa Fluor 488 and Alexa Fluor 594 were diluted in 50 mM sodium phosphate buffer, pH 7 with 50 mM NaCl and mixed together to the final concentration such that optimal amount of labels were diffusing through confocal volume at the same time. The relative cross-correlation of free dyes was measured. The amplitude of negative control cross-correlation function G (τ) is 1.032, which indicates that only 3.2% of signal is caused by spectral cross-talk and nonspecific binding of fluorophores. The positive control - fluorescently labeled complementary DNA oligonucleotides were hybridized and measured under the same conditions as the negative control.

**Figure 2-figure supplement 2**. Microscale thermophoresis (MST) assays. (**a**) TRF1-TIN2 interaction measurements show K_D_ 240 nM. (**b**) TRF2-TIN2 interaction displays higher K_D_, hence lower affinity of TRF2 to TIN2. MST assay conditions are described in Material and Methods.

**Figure 2-figure supplement 3**. 10%SDS PAGE gels of purified proteins. From left: TRF1, TRF2, TIN2 and TPP1. Gels were stained by Bio-Safe Coomassie G-250.

**Figure 4-figure supplement 1**. The animation of gradual addition of TRF2 to the solution of 10 nM fluorescently labeled telomeric DNA R5 induces aggregates that are visible by fluorescence microscopy. Experimental conditions are the same as described in Figure 4. The aggregates were formed also when TRF2 lacking the basic B-domain was used.

**Figure 5-figure supplement 1**. TPP1 is essential for TRF1-TIN2-TPP1-TRF2 complex assembly. Relative cross-correlation of fluorescently labeled TRF1 (Alexa Fluor 594) and TRF2 (Alexa Fluor 488) was measured upon addition of unlabeled TIN2 or preformed TIN2-TPP1 complex. The relative cross-correlation corresponds to the proportion of cross-correlation amplitude related to the TRF2 auto-correlation amplitude. The low relative cross-correlation indicates no interaction between TRF1 and TRF2, hence TIN2 was insufficient to interconnect TRF1 and TRF2 into a stable complex. On the contrary, upon addition of preformed TIN2-TPP1 complex, the significant increase of relative cross-correlation of TRF1 and TRF2 was observed. The relative cross-correlation increase demonstrates that the TRF1 and TRF2 were simultaneously diffusing through the confocal volume together. The TRF1-TIN2-TPP1-TRF2 complex was formed only if TPP1 was present.

**Figure 5-figure supplement 2**. A110R and A15R mutations of TIN2 inhibit shelterin core complex formation. (a) Alexa Fluor 488 labeled TIN2-A110R and unlabeled TRF1 were incubated to form a complex and solely Alexa Fluor 594 labeled TRF2 or mixture of labeled TRF2 with unlabeled TPP1 was added. The amplitude of relative cross-correlation between TRF2 and TIN2-A110R remained low which means that the shelterin-core was not formed. After replacing the TIN2 mutant variant for TIN2, the relative cross-correlation significantly increased, hence the shelterin core was formed. (b) The relative cross-correlation of Alexa Fluor 488 labeled TIN2-A15R and Alexa Fluor 594 labeled TRF2 was measured after adding of TRF2 or mixture of TRF2 and unlabeled TPP1 of the preformed mixture of TIN2-A15R and unlabeled TRF1. The amplitude of relative cross-correlation remained low, hence the shelterin core was not formed.

